# *TP53* and *RB1* are predictive genetic biomarkers for sensitivity to cytarabine in gliomas

**DOI:** 10.1101/2025.11.20.688938

**Authors:** Liisi Laaniste, Joachim Luginbuehl, Mi Trinh, Rachel Watkins, Gregor Lueg, Grant Neilson, Marjolein Burgers, Kasia Kania, Regina Reynolds, Zoe Smith, Julia Kleniuk, Lauren Gould, Dorota Pregowska, Sophie Mathiesen, Michael Johnson, Louwai Muhammed

## Abstract

Therapeutic progress in glioma, one of the most lethal human cancers, has been limited by molecular heterogeneity and lack of biomarker-driven drug deployment. Here we used a proprietary large-scale CRISPRi screening in primary patient-derived glioma tumorspheres to identify genetic vulnerabilities and nominate pharmacologically tractable targets. DNA polymerase-linked dependencies emerged as top-ranked hits, which we validated through orthogonal viability assays. Network-based integration of dependency data with drug-target relationships nominated cytarabine, a nucleoside analogue already approved for intrathecal use, as a candidate agent targeting this axis. Dose-response profiling across molecularly diverse glioma models revealed substantial heterogeneity in cytarabine sensitivity (IC_50_ range: 0.04-9.8 µM). Machine learning analysis of whole-genome sequencing data identified *TP53* wild-type and *RB1*-wild-type status as dominant predictors of response, with double wild-type lines showing three standard deviations (3 s.d.) increased sensitivity compared to altered models. Prospective validation in an independent cohort confirmed that *TP53/RB1* genotype stratifies cytarabine activity. These findings establish a mechanistically anchored, biomarker-restricted repurposing opportunity for cytarabine in leptomeningeal glioma, enabling rational prioritisation of an accessible therapy in a molecularly defined patient subset.

## Introduction

Gliomas are primary brain tumours and remain among the most lethal human cancers. Despite major advances in molecular characterization over the last decade, therapeutic progress remains modest. Historically, most interventional trials in glioma have been conducted without molecular stratification, diluting an efficacy signal in genetically heterogeneous populations and likely contributing to the high rate of trial failure. Temozolomide (TMZ), the backbone of standard-of-care treatment for glioblastoma multiforme, achieves benefit primarily in tumours with *MGMT* promoter methylation, illustrating that even established agents operate within specific genetic contexts [1]. More recently, the IDH inhibitor Vorasidenib and the DRD2 antagonist ONC-201 have demonstrated activity in biomarker-defined populations of glioma namely, *IDH*-mutant glioma and *H3K27M*-mutant diffuse midline glioma, respectively [2], [3]. These examples underscore the principle that efficacy in glioma is frequently contingent on underlying genetic context. Identification and prospective deployment of genetic biomarkers to match drugs to molecularly defined subgroups are therefore fundamental for rational drug development in glioma.

Systematic functional genomic screening provides a direct route to map genotype-linked vulnerabilities that are not always apparent from genomic annotation of mutations and copy number alterations alone. By perturbing thousands of genes across large panels of molecularly profiled tumour models, it is possible to resolve context-specific dependencies and nominate biomarkers that can both explain and predict drug response. Integrating perturbation data with somatic alterations has consistently revealed that genetic aberrations can generate collateral dependencies susceptible to pharmacological targeting [4], [5], [6]. These resulting maps of induced dependency provide a mechanistic framework for aligning drugs with the molecular contexts that determine their efficacy, enabling a rational, rather than empirical, approach to therapy in genetically heterogeneous cancers such as glioma.

Here we used dependency scores from a large functional genomic screen in primary, patient-derived glioblastoma tumour spheres (unpublished) to identify selective, genetically-conditioned vulnerabilities. We first nominated genes whose perturbation produced strong loss-of-viability in a select population of samples. We then embedded these lethal hits into a functional interaction network to resolve convergent pathways and mechanistic nexuses that could be pharmacologically intercepted. This analysis surfaced a tractable axis targeted by the clinically available drug cytarabine. We next asked whether cytarabine sensitivity was genetically stratifiable and identified *TP53* and *RB1* alterations as candidate biomarkers of response. Finally, we prospectively validated both the predicted efficacy and its biomarker restriction in an independent cohort of newly established glioma tumour spheres, supporting a mechanistically anchored, biomarker-driven repurposing opportunity for this agent in a molecularly defined glioma subset.

These findings have potential immediate clinical applications, as cytarabine is already approved for intrathecal administration in acute myeloid leukaemia (AML) with leptomeningeal involvement and is used off-label for leptomeningeal dissemination of glioma [7], [8], [9], [10]. Our findings suggest that, in patients whose tumours harbour the relevant biomarker genotype, cytarabine could be rationally prioritised over other intrathecal agents currently used empirically in this setting, enabling a genomically-informed redeployment of an accessible therapy.

## Results

### Polymerase-linked dependencies emerge as top-ranked liabilities and nominate cytarabine as a tractable axis for pharmacologic interception

To identify genetic vulnerabilities in glioblastoma we made use of functional genomic screening data from pooled CRISPRi screen containing 57,624 sgRNAs targeting 12,832 protein-coding genes in 26 primary, patient-derived glioma tumour spheres and 2 neuroprogenitor spheres acting as a control (unpublished data). Positively or negatively selected genes were determined using the MAGeCK algorithm [11]. We constructed a ranked vulnerability landscape by scoring gene-level effects using MAGeCK-RRA. We combined the adjusted p-values per gene across all samples using the Simes method. Pathway enrichment analysis of significantly (adjusted combined p-value of < 0.05) depleted genes (n = 615) indicated the strongest pathway level dependency in DNA replication (**Fig. 1s**). Notably, genes converging on the DNA Pol-α/δ/ε replication apparatus clustered within the highest-ranking end of the distribution, indicating dependencies in DNA metabolism–linked replication and transcriptional programmes (**Fig. 1a**).

**Fig. 1.**
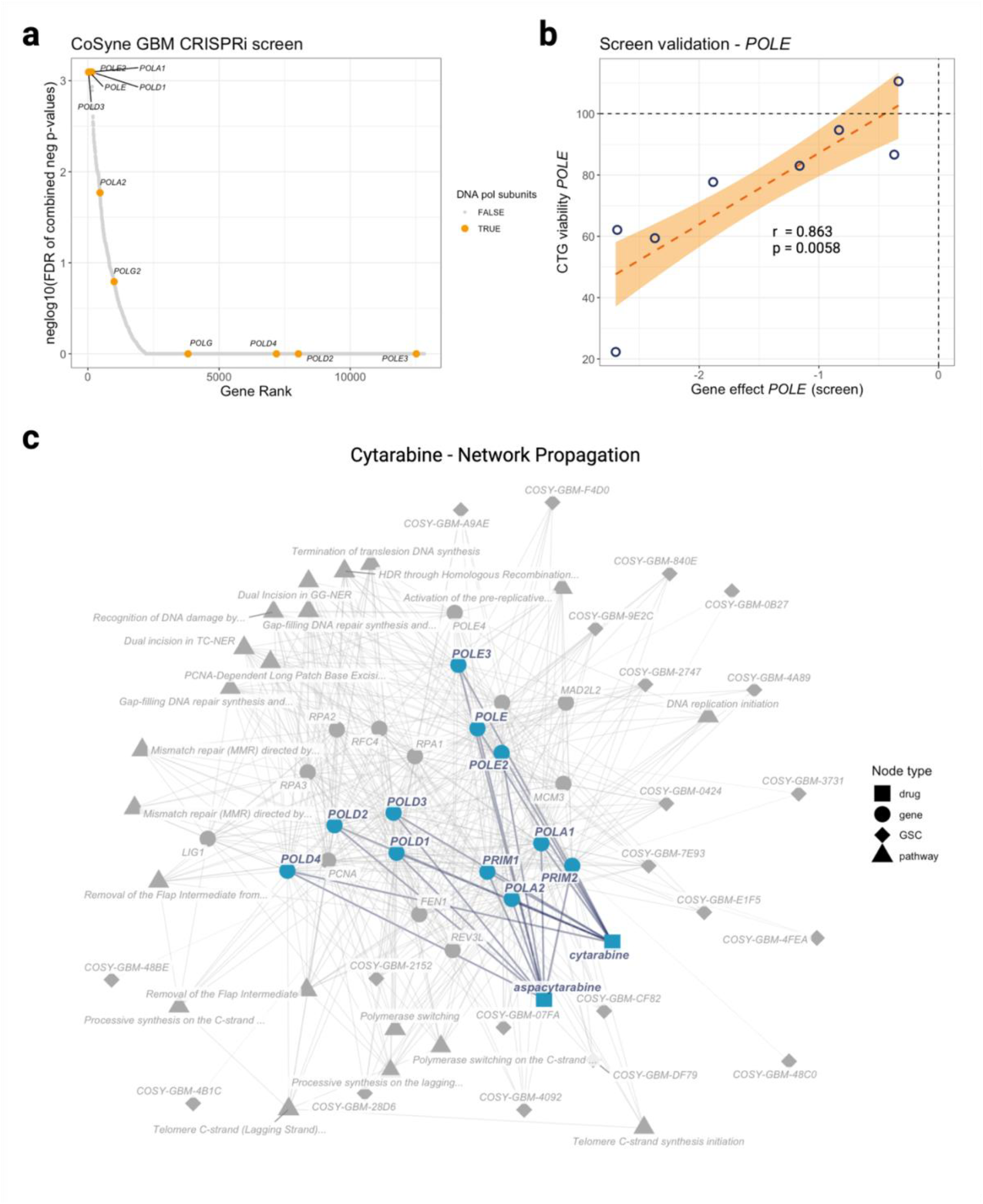
**a**, Ranked gene-level dependencies from primary patient-derived pooled CRISPRi screen in 3D glioma, based on a ranking of combined adjusted p-values with the Simes method. Selected genes with known function in DNA polymerase complex. **b**, Validation of *POLE* dependency using targeted CRISPRi and a direct cell-viability assay (CellTiter-Glo 3D) in 3D glioma spheres (*n* = 8). Each point represents the mean viability from three independent biological replicates, normalized to non-targeting control and positive control Paclitaxel. Shaded bands indicate 95 % confidence intervals around the fitted regression line. **c**, Network-based nomination of cytarabine as a compound targeting the polymerase-linked vulnerability cluster. A graph-based integration of CRISPRi dependencies with pathway annotations, molecular functions and drug–target relationships (subgraph based on the insights graph with 21,374 nodes and 1,234,908 edges) revealed a dense convergence of high-ranking hits on the DNA polymerase replication apparatus (blue nodes). Cytarabine and its pro-drug aspacytarabine (blue squares) were identified as compounds with annotated activity against this axis. Edges indicate curated relationships among genes, processes, genetic dependencies and drugs.

We selected one of the highest-ranking genes of the complex, *POLE* (DNA polymerase-ε, leading-strand replication), as a representative node within the polymerase-linked vulnerability cluster. To orthogonally test whether the CRISPRi screening signal reflected true lethality, we inactivated *POLE* using targeted CRISPRi in matched 3D patient-derived glioma spheres (n = 8) and quantified cell viability using a CellTiter-Glo (CTG) assay (**Fig. 1b**). CRISPRi dependency scores for *POLE* significantly correlated with the degree of viability loss in the CTG assay (Pearson r = 0.863, p = 0.0058), indicating that CRISPRi screen-derived essentiality calls reproducibly translate into direct cell-death phenotypes in primary 3D glioma cultures.

The strength and coherence of this polymerase-associated signal provided a rationale to investigate whether this axis could be pharmacologically intercepted and genetically stratified. We therefore generated a subgraph based on a proprietary empirical insights graph (total size of the insights graph: 280,678 nodes and 21,374,777 edges). This subgraph integrates primary 3D patient derived glioma CRISPRi screen dependency calls with curated pathway annotations, biological process ontologies, molecular functions, and drug–target relationships to identify compounds with activity against polymerase-linked mechanisms (see method). To identify drugs that hit genes (and pathways) that show strong dependency in the screen, we leveraged a constrained PageRank algorithm over a heterogeneous subgraph with the start node being the represented drug nodes [12]. This analysis nominated cytarabine and its pro-drug aspacytarabine as candidates targeting this axis (**Fig. 1c**).

### Cytarabine displays biomarker-stratified activity in primary patient-derived glioma models defined by TP53/RB1 wild-type genotype

To translate the computational nomination into experimental evidence of drug effect, we set out to profile cytarabine dose-response in a diverse set of glioma cell lines. To capture cellular heterogeneity, we applied principal component analysis on bulk RNA-seq data from 48 glioma tumour spheres and selected 15 cell-lines along PC1 for the drug assays (**Fig. 2a**). CTG assay of cytarabine response displayed a wide dynamic range of activity, with IC_50_ values spanning nearly 2.4 orders of magnitude (0.04–9.8 μM), with an average IC_50_ of 2.2 μM, indicating substantial intrinsic heterogeneity in susceptibility to this agent across primary glioma models (**Fig. 2b**).

**Fig. 2.**
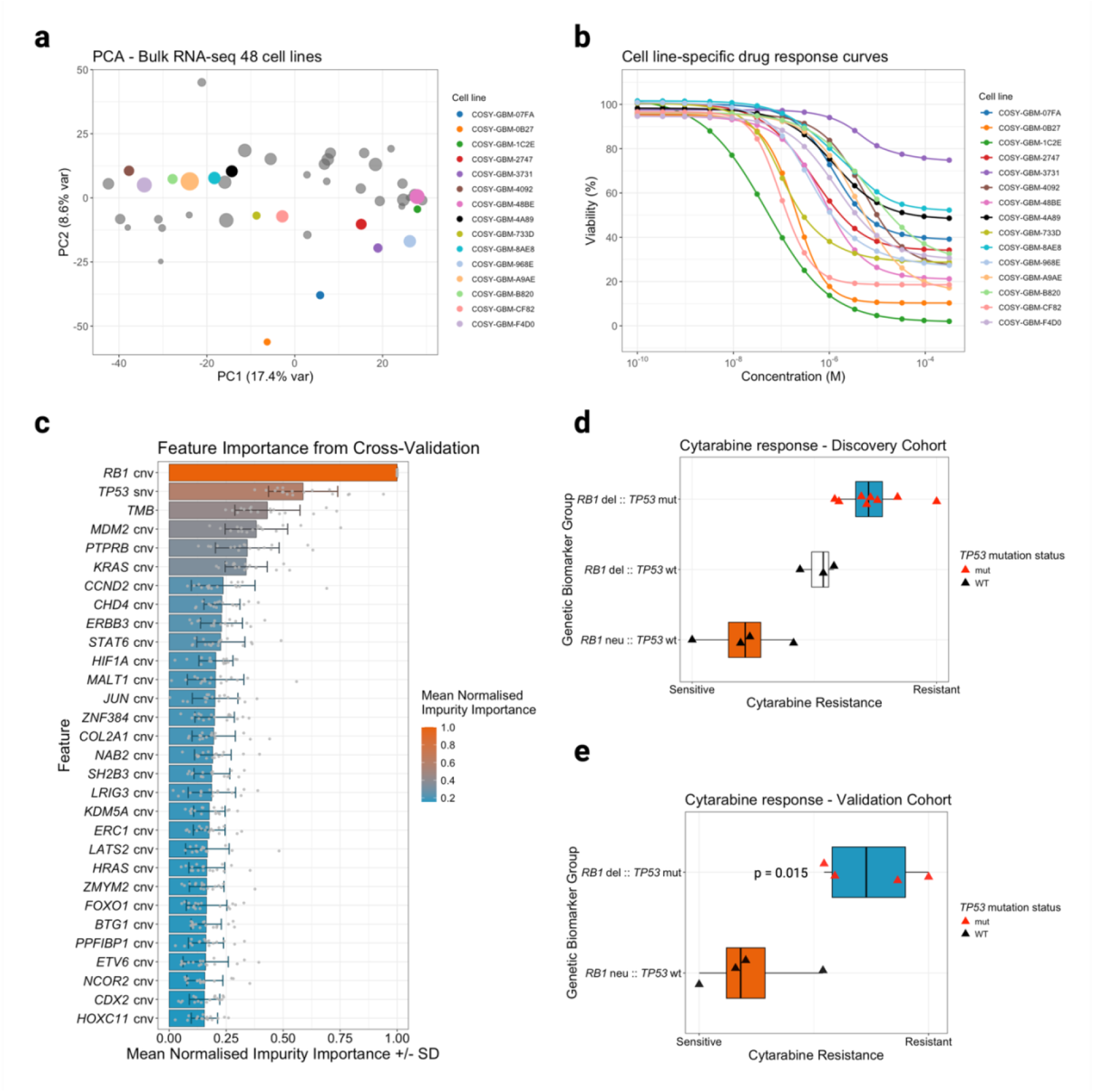
**a**, Principal component analysis of transcriptomic profiles from 48 patient-derived GBM cell lines. The 15 lines selected for cytarabine dose–response assays are highlighted in colour. Point size corresponds to cell-doubling time, with smaller points indicating faster-growing lines. The first principal component (PC1) captures the major axis of transcriptional heterogeneity (17.4% variance) used for representative line selection. **b**, Cytarabine dose–response curves across patient-derived 3D glioma tumour spheres (*n* = 15). Cells were treated with cytarabine across a 14-point concentration range (10^−4^–10^−9^ M) for 6 days and viability was measured using the CellTiter-Glo 3D assay. Each curve represents the mean of three independent biological replicates per concentration, normalized to DMSO controls. IC_50_ values spanned ∼2.4 orders of magnitude (0.04–9.8 µM), indicating substantial heterogeneity in cytarabine sensitivity among primary GBM models. **c**, Bar plot showing the top 30 genetic features predicting cytarabine response, ranked by their mean impurity-based importance scores from the random-forest model. Scores were normalised to a 0-1 scale; error bars indicate ±1 standard deviation across model runs. Features with higher importance contributed more strongly to separating sensitive from less sensitive cell-lines to cytarabine. TMB - Tumour mutation burden. **d**, *TP53/RB1* genotype stratifies cytarabine response in patient-derived glioma models. Normalised cytarabine viability scores (aggregated across drug doses) are shown for glioma lines grouped by *RB1* and *TP53* status. *RB1*-neutral / *TP53*-wt lines (orange) exhibited the lowest viability (greatest sensitivity), *RB1*-deleted / *TP53*-wt lines (grey) showed intermediate sensitivity, and *RB1*-deleted / TP53-mutant lines (blue) were least sensitive (highest viability). Individual lines are overlaid as points (triangles denote *TP53* status). This pattern indicates that intact *TP53* and *RB1* pathways are associated with heightened cytarabine susceptibility. **d**, Mean log_10_(IC_50_) values for cytarabine are shown for glioma lines classified as responders (*TP53/RB1*-neutral; blue) versus non-responders (*TP53* and/or *RB1*-altered; red). Responder lines exhibit markedly lower IC50 values, consistent with greater cytarabine sensitivity. Points represent individual cell lines; boxes show interquartile range with median. **e**, Four TP53/RB1 wild-type and four *TP53/RB1*-altered lines were treated with cytarabine across a 14-point concentration range (10^−4^–10^−9^ M). Viability was quantified by CellTiter-Glo 3D after 9 days and normalized to DMSO controls. Data represent the mean of three independent biological replicates per cell line. Box plots show interquartile range with median; individual points represent the mean of 3 biological replicates. *TP53/RB1*-neutral lines exhibited significantly lower normalized viability, consistent with enhanced cytarabine sensitivity (*p* < 0.05, two-sided t-test).

We hypothesized that genetic diversity of patient-derived glioma tumour spheres could underpin the observed differential drug response. To identify a genetic biomarker of response, we leveraged whole-genome sequencing data from the assayed tumour spheres, specifically single nucleotide and copy number variants, and applied a random-forest regression model with feature-importance ranking to nominate genetic alterations predictive of cytarabine sensitivity (measured by the area under the dose-response curve or AUC). *RB1* and *TP53* alteration status emerged as the dominant features predicting the difference in cell viability after drug treatment, indicating that wild-type status of these tumour suppressors defines a biomarker-linked sub-population with heightened vulnerability to cytarabine (**Fig 2c**).

Glioma cell lines with *TP53* wild-type and *RB1* copy-number neutral status substantially greater responsiveness to cytarabine compared to *TP53*-mutant and *RB1*-deleted models, corresponding to a mean AUC difference of approximately three pooled standard deviations between groups (Hedges’ g = 3.34, 95% CI 1.49 - 5.13). Notably, *TP53* wild-type in an *RB1*-deleted background still conferred higher sensitivity relative to double-altered (*TP53*-mutant/*RB1*-deleted) models (Hedges’ g = 1.64, 95% CI 0.19 - 3.04), indicating independent and accumulative influence of these genetic alterations on cytarabine response (**Fig. 2d**).

To validate the genetic biomarker prediction, we tested prospectively an independent validation set of eight additional patient-derived cell lines not included in the discovery cohort (n = 4 *TP53/RB1* wild-type and n = 4 with *TP53/RB1* alterations). Cells were treated with cytarabine for 6 days across a 14-point serial dilution range (10^−4^ to 10^−9^ M) and viability was measured by CTG assay. Consistent with the model-derived prediction, *TP53*/*RB1*-wild-type lines were significantly more sensitive than altered lines (p = 0.016, Welch’s t-test), confirming that *TP53/RB1* genotype stratifies cytarabine response in glioma (**Fig. 2e**).

## Discussion

Genetic dependency mapping enables systematic quantification of gene importance, as conditioned by the underlying genetic context of cancer cells. Such genetic perturbation data can be used to identify mechanistically linked novel therapeutic targets, and to reveal genetic biomarkers that stratify the activity of existing clinical stage compounds. This provides a rational basis for precision deployment of cancer drugs to the patient subgroups most likely to respond. The present work demonstrates that a polymerase-linked dependency signal emerging from a large-scale functional genomic screen in primary glioma 3D tumour spheres is reproducible across orthogonal assays, and identifies a genetically stratified, pharmacologically actionable node anchored on an already-approved agent. By integrating CRISPRi-derived essentiality with network-based target mapping, drug-target annotation, and *in vitro* validation across molecularly diverse patient-derived models, we show that cytarabine, already in clinical use for CNS disease, displays pronounced and genotype-linked activity in glioma. In this setting, *TP53*/*RB1* genetic alteration status emerged as the dominant predictor of sensitivity.

Cytarabine has long been used to treat AML, including leptomeningeal metastasis of AML, where *TP53* mutation has been associated with differential cytarabine responsiveness [13]. This supports the idea that p53 status can modulate nucleoside-analogue efficacy in human disease contexts. Our findings extend this concept to glioma and suggest that *TP53* wild-type status similarly conditions heightened cytarabine sensitivity in primary glioma models, albeit in an entirely different lineage. In addition, *RB1* emerged as a second biomarker of cytarabine response in glioma, to our knowledge not previously reported in this context and not a recognised determinant of cytarabine response in AML, indicating that the biomarker logic of cytarabine may be tissue-specific. Notably, *CDKN2A/B* deletion, a canonical cell-cycle lesion in glioma and a known modulator of cytarabine responses in AML, did not contribute significantly towards stratifying cytarabine sensitivity (**Fig. 2c**) [14]. This underscores the non-transferability of AML-derived assumptions to gliomas and reinforces the need to derive biomarker frameworks directly from glioma biology, rather than externally importing them.

In glioma, intrathecal cytarabine has been used off-label in leptomeningeal dissemination of the disease on an empiric basis in selected patients, with case reports documenting durable symptomatic and radiographic responses in those individuals [8], [9], [10]. In an in vitro 3D glioma tumour sphere setting, the average IC_50_ of cytarabine was 2.2 μM. Pharmacokinetic studies of intrathecal cytarabine of a single dose of 100 mg at 48-h intervals show mean peak cytarabine concentration of 16.7 ± 6.3 mM and a mean concentration of 0.77 ± 0.53 mM after 6 h. IC_50_ concentrations observed in the in vitro assay are therefore in a clinically relevant range [15].

The identification of *TP53* wild-type and *RB1*-neutral states as predictors of heightened cytarabine susceptibility in primary glioma models provides a mechanistic and testable rationale to move from unstratified intrathecal deployment to biomarker-enriched use in leptomeningeal disease. Prospective clinical stratification of patients with leptomeningeal glioma by *TP53/RB1* status could therefore enrich for responders, reduce futile exposure in predicted non-responders, and help clarify whether the previously observed responders in case literature is in fact a molecularly coherent group rather than a clinical anomaly. Given that intrathecal cytarabine is already clinically deliverable within current practice guidelines for leptomeningeal metastases, these observations support an immediately feasible, biomarker-anchored refinement of an existing therapy rather than the introduction of a new agent.

Together, these data argue that cytarabine is not merely an empirically active agent, but a genetically-stratifiable therapeutic opportunity in leptomeningeal glioma. *TP53/RB1*-based enrichment could rationally guide its prioritisation over other intrathecal or systemically delivered agents already considered in the neuro-oncology setting for patients suitable to receive such treatment.

## Methods

### Data availability

The functional genomic screening data used to generate the hypothesis in this study are not openly available due to reasons of sensitivity, and are available from the corresponding author upon reasonable request.

### Generation of a single ranked gene list

To combine the results across the 26 primary, patient-derived glioma tumour spheres (unpublished data), the negative p-values from the MAGeCK RRA algorithm were combined with the Simes method [16]. Briefly, for each gene, the p-values across all cell lines were sorted in ascending order, and the combined p-value was calculated as min (n × p_(i) / i), where n is the total number of cell lines and p_(i) is the i-th smallest p-value. This approach accounts for dependencies between tests whilst controlling the family-wise error rate under certain dependence structures. The combined p-values were subsequently adjusted for multiple testing using the Benjamini-Hochberg procedure to control the false discovery rate. Lower adjusted p-values indicate stronger evidence for consistent negative selection of the gene across multiple cell lines.

### Over-enrichment analysis of depleted genes

Genes with an adjusted p-value ≤ 0.05 (N = 615) from the Simes method were subjected to over-representation analysis using a hypergeometric test against Reactome pathways [17]. The gene universe was defined as all genes included in the CRISPRi screen. Enrichment p-values were adjusted for multiple testing using the Benjamini-Hochberg procedure, and pathways reaching an FDR threshold of ≤ 0.05 were considered significantly enriched. To visualise relationships between enriched pathways, an enrichment map [18] was constructed where nodes represent individual pathways and edges connect pathway pairs with a Jaccard similarity coefficient of at least 0.25. The colour coding of the nodes represents the average of all the - log_10_(combined FDR) from the Simes method for the genes in this pathway. Darker colouring indicates that on average, more genes are present in the pathway that show consistent negative selection.

### Empirical Data Enhanced Insights Graph

A proprietary graph-based database was constructed integrating various public resources including OpenTargets [19], Reactome [20], STRING [21], Signor [22], Gene Ontology [23], Interpro [24], and more, as well as internally derived edges from analysis of the biobank. The total size of the graph database is 280,678 nodes and 21,374,777 edges.

### Network-based identification of cytarabine and its pro-drug aspacytarabine

A subgraph was constructed from the insights graph comprising the following components:

Nodes: 28 primary patient-derived gliomas from the CRISPRi screen; 18,309 genes/proteins represented in the screen or network resources; 2,751 Reactome pathways; and 286 drugs selected based on being approved for human use and targeting genes significantly depleted (adjusted p-value ≤ 0.05) in at least one glioma line.

Edges: 556,360 protein-protein interaction edges derived from STRING [21] (interaction score ≥ 0.85), IntAct [25] (physical associations), Reactome [20] protein-protein interactions and SIGNOR [22], merged and deduplicated; 5,538 hierarchical edges connecting Reactome pathways; 46,585 edges linking Reactome pathways to their constituent genes/proteins; 14,368 edges connecting glioma lines to their significantly depleted genes (adjusted p-value ≤ 0.05 from MAGeCK); and 1,787 drug-target edges. In all cases, reverse edges were added to the graph, effectively doubling the total edge count.

A constrained PageRank algorithm [26] was applied using each of the 286 drugs individually as the starting node in the personalisation vector. The random walk was configured such that upon reaching a glioma node or another drug node, the walker was forced to restart at the initial drug node. This approach quantified each drug’s influence across the network. As DNA replication and polymerase machinery emerged as the most significantly depleted biology from the Simes combined p-value analysis, drugs with the highest influence on these pathways were prioritised, identifying cytarabine and its pro-drug aspacytarabine as lead candidates.

### Principal component analysis of transcriptomic diversity

Principal component analysis was performed on bulk RNA-seq data (unpublished) from 48 patient-derived GBM tumour sphere lines to capture global transcriptional heterogeneity. Raw count data were log_1#x002B;_-transformed and the 3,000 most variable genes across samples were selected. Gene identifiers were sanitized by removing Ensembl version suffixes and transcript annotations. Expression values were centred and scaled per gene, and principal component analysis was computed using the prcomp function in R (v4.3.2).

### CRISPRi-mediated POLE knock-down and CellTiter-Glo viability assay

To validate gene-level dependencies identified in the pooled CRISPRi screen (Fig. 1a), individual knock-downs were performed in eight patient-derived GBM tumour sphere lines stably expressing dCas9–KRAB. Three sgRNAs previously included in the pooled screen targeting *POLE* were cloned under a U6 promoter into lentiviral vectors (EF1α–BSD–T2A– EGFP backbone). Equal amounts of the three plasmids were pooled for virus production in HEK-293 FT cells. Briefly, HEK-293 FT cells were transfected at 60–70% confluence with the CRISPRi expression vector and packaging plasmids (psPAX2 and pMD2.G) using PEIpro (Polyplus #101000017) in Opti-MEM. Viral supernatants were collected at 48 h and 72 h, clarified, PEG-precipitated, and resuspended in PBS before storage at –80 °C. For the viability assay, glioma spheroids were seeded in 96-well ultra-low-attachment (ULA) black-walled plates at 2,000 cells per well in 100 µL of complete NeuroCult NS-A medium (STEMCELL Technologies #05750/05753) supplemented with heparin, EGF, FGF, and Pen/Strep. Cells were infected the following day with pooled lentivirus at a multiplicity of infection (MOI) of 2 to ensure that each cell received at least one sgRNA. Polybrene (8 µg/mL) was added during infection. A non-targeting sgRNA control virus was included in parallel to serve as a negative control for cell viability. After centrifugation (100 × g, 3 min) and 7 days of incubation, cell viability was quantified using the CellTiter-Glo 3D Cell Viability Assay (Promega #G9683) according to the manufacturer’s instructions: 100 µL reagent per well, shaking 5 min, room-temperature lysis 25 min, and luminescence read-out on a CLARIOstar plate reader. Viability values were normalized to the non-targeting control wells. Parallel samples from each cell line were collected for RNA extraction and knock-down quantification using the TaqMan™ Fast Advanced Cells-to-CT™ Kit (Invitrogen #A35374, A35377, A35378) with gene-specific assays for *POLE* and *ACTB*.

### Drug treatment and viability assay

To assess pharmacologic sensitivity, patient-derived 3D GBM tumour spheres were treated with cytarabine (Merck #C1768) across a 14-point serial dilution range (10^−4^ to 10^−9^ M, 1:3.16 steps) in complete NeuroCult NS-A medium containing ≤ 0.1 % DMSO. Cells were seeded at 2,000 cells per well in 96-well ultra-low-attachment (ULA) plates and allowed to form spheroids for 72 h prior to treatment. The drug was applied in two sequential 72-h doses (Day 3 and Day 6) using freshly prepared dilutions. Puromycin was included as a positive (lethal) control and DMSO as vehicle control. At Day 9, cell viability was quantified using the CellTiter-Glo 3D assay as described above for the CRISPRi validation experiments. Luminescence values were background-subtracted, normalized to DMSO controls, and dose– response curves were fitted using a four-parameter logistic model in R to derive IC_50_ and AUC (area under the dose-response curve) values. For biomarker validation (Fig. 2E), an independent cohort of eight glioma tumour sphere lines (four *TP53/RB1* wild-type and four with *TP53* and *RB1* alterations) were treated with the same cytarabine concentration range under identical assay conditions, and viability was measured as above.

### Cytarabine-response biomarker discovery with random forest

To identify potential genetic alterations predictive of cytarabine sensitivity, we trained a random forest regression using AUC as a quantitative measure of drug response. Lower AUC values indicate greater sensitivity (viability loss at lower drug concentrations), whereas higher AUC values indicate relative resistance. Random forest modelling was performed using the ranger R package.

Genetic features were encoded as binary indicators of gene-level aberrations (somatic mutations and copy-number changes) per cell-line and restricted to Tier 1 genes in the COSMIC Cancer Gene Census (v101) to focus on cancer-relevant genes. Tumour mutation burden (TMB) and estimated doubling time for each cell-line were also included as features. In total, 528 features were provided as inputs for model training.

Model performance was assessed using leave-one-out cross-validation (n = 15), with prediction accuracy evaluated by root mean squared error and Pearson correlation between predicted and observed AUC values.

Feature importance was quantified by mean decrease in node impurity. Importance scores were min-max normalised to scale between 0 and 1 within each cross-validation iteration and averaged across iterations to assess stability. SHAP (SHapley Additive exPlanations) values were also computed to quantify each feature’s contribution to the predicted AUC across individual cell lines, using the TreeSHAP algorithm. SHAP scores capture both the direction and magnitude of a feature’s effect, providing insights into whether a genetic event increases or decreases predicted drug sensitivity. For each cross-validation iteration, SHAP scores were scaled by the maximum absolute SHAP value within that iteration, resulting in scores ranging from-1 to 1 (Fig. 2s).

Relative cytarabine sensitivity between biomarker-stratified groups was quantified as the percent decrease in group-mean *AUC*: 100 × (*AUC*_*A*_ − *AUC*_*B*_)/*AUC*_*A*_

All human glioma samples used in this study were collected with informed consent and in accordance with institutional ethical guidelines.

## Supplementary Figures

**Fig. 1s.**
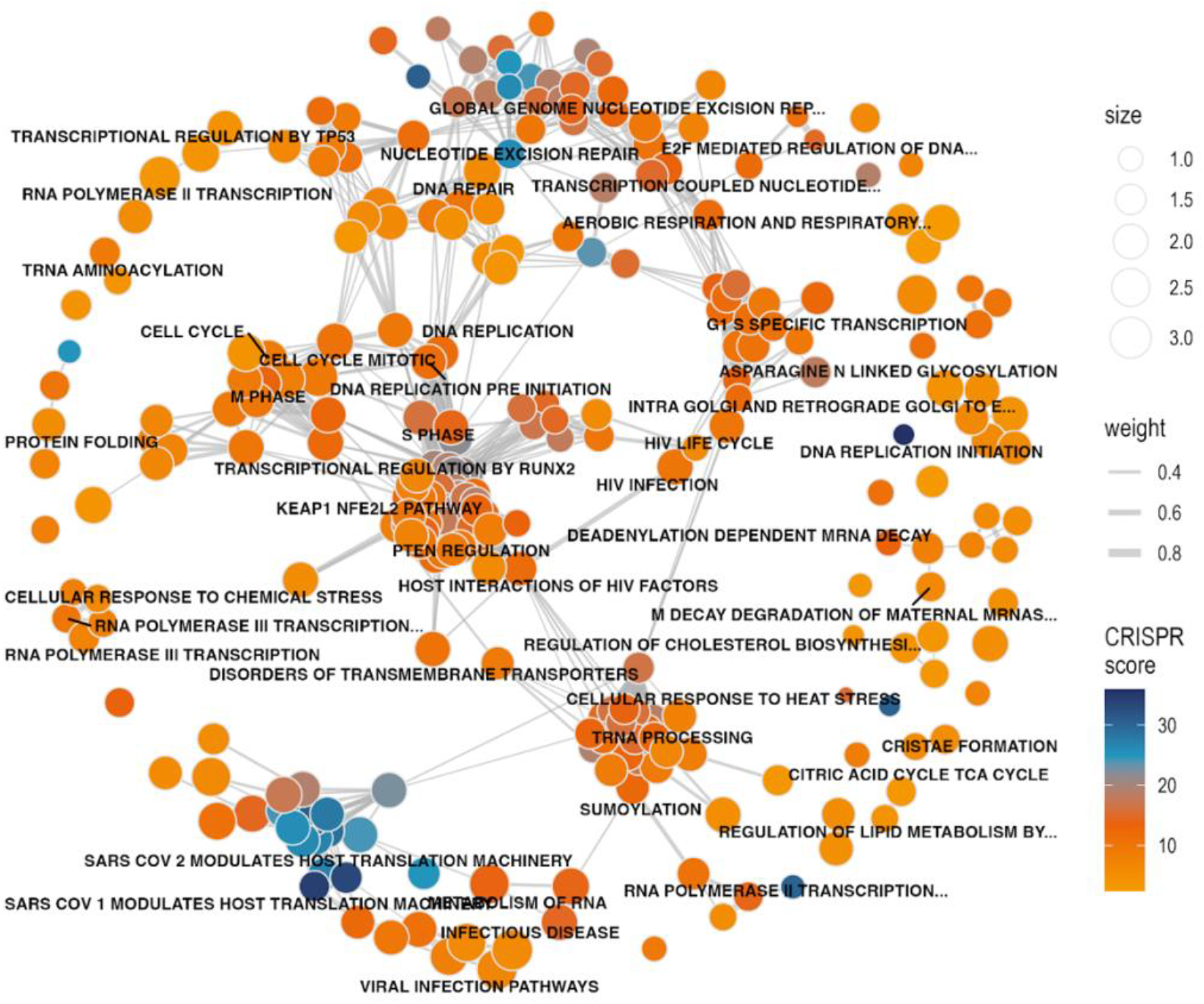
Enrichment map of Reactome pathways of all genes that showed significant combined depletion (after FDR correction) by the Simes method (N = 615). The edge weight (Jaccard similarity) between two pathways had to be at least 0.25. Node colour (CRISPR score) represents the average-log10(Simes combined FDR) and indicates how dependent the screened GBM cell lines are on this particular pathway. Dark blue (for example, DNA replication initiation) indicates strong, significant dependency to member genes in this given pathway.

**Fig. 2s.**
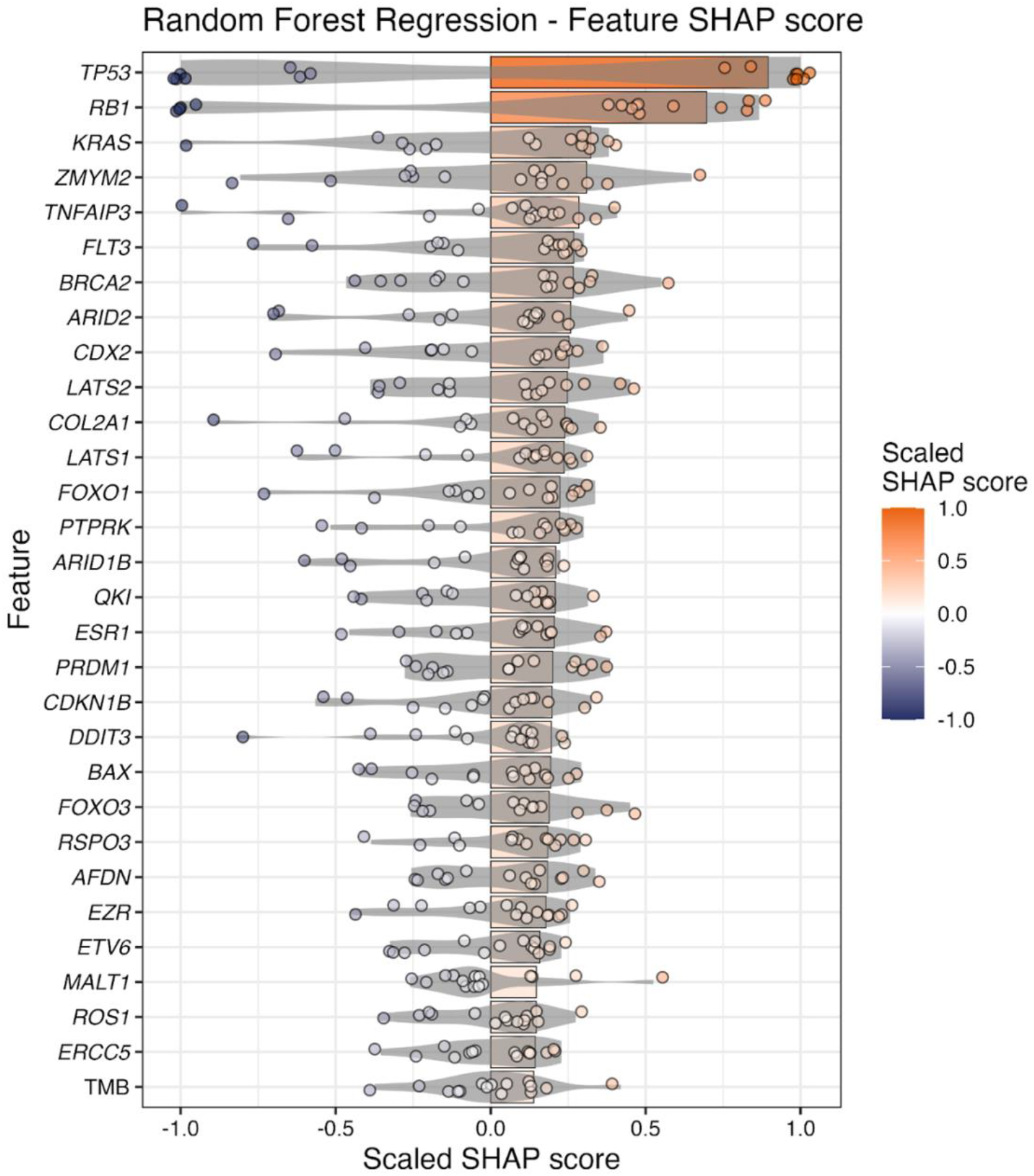
SHAP scores for top 30 genetic alterations predicting cytarabine response. Bar plot showing the mean absolute SHAP scores of the top 30 genetic alterations across leave-one-out cross-validation iterations. For each iteration, SHAP scores were scaled by the maximum absolute SHAP value, resulting in scores ranging from-1 to 1. For each feature (y-axis), individual dots represent the scaled SHAP score of that feature for the left-out sample. Features with high SHAP variance exhibit more polarised or bimodal effects, indicating stronger discriminatory power between cytarabine-sensitive and-resistant cell lines.

